# Targeting granule initiation and amyloplast structure to create giant starch granules in wheat

**DOI:** 10.64898/2025.12.12.693964

**Authors:** Rose McNelly, Lara Esch, Qi Yang Ngai, Katalin Pohan, Rhea Stringer, Brendan Fahy, Frederick J Warren, David Seung

## Abstract

Starch granule size influences the functional, digestive, and processing qualities of starch, but its genetic control in plants is poorly understood. Here, we demonstrate that both space and substrate constraints limit starch granule size in wheat and provide an approach to achieve substantial increases in granule size. Wheat starch typically contains large A-type and small B-type granules. To increase A-type granule size, we explored the effect of mutations in the plastid division component *PARALOG OF ARC 6* (*PARC6*), which increases amyloplast size and therefore the space available for granule growth, and in *B-GRANULE CONTENT 1* (*BGC1)*, which reduces the number of granule initiations and thus competition between growing granules for space and substrates. While *parc6* and *bgc1* single mutants had only modest increases in A-type granule size, the *parc6 bgc1* double mutant produced striking giant granules that were more than double the size of typical A-type granules. The increase in granule size in *parc6 bgc1* was reproducible in both the glasshouse and field, and had no detectable effect on plant growth, grain size and starch composition or content. We demonstrate that the size increase affects a range of functional properties, including viscosity and pasting temperature. Overall, by targeting both constraints, we created a new class of giant cereal starch that has not been previously observed in nature, with altered physicochemical properties that can be used in food and industrial applications.

**Significance statement:** We provide a major advance in understanding the factors determining starch granule size in plants – a trait that strongly influences starch functionality. Starch granules from cereals are typically smaller than those of most root/tuber crops, rarely exceeding 30 µm. We created a novel cereal starch in wheat that exceeds this size range. By simultaneously increasing amyloplast size and reducing granule initiations, we increased space for granule growth and reduced competition between growing granules for substrates. This resulted in a new class of cereal starch where nearly 50% of the starch volume were in granules greater than 30 µm, and some exceeded 50 µm. These giant granules had altered physicochemical properties that could find novel application in food and industry.

## Introduction

Starch is a fundamentally important resource – it contributes up to 50% of our dietary calories, and is used in the industrial production of paper, textiles, adhesives, biofuels and biopolymers (1). It is synthesised by plants and comprised of amylose and amylopectin, which are glucose polymers. These polymers assemble into insoluble, semi crystalline starch granules in plastids (2, 3). There is large interspecies variation in storage starch granule morphology in terms of granule shape and size (4, 5). Cereal starch granules typically range from 2-30 µm, and are smaller than those found in other crops including pea (10-45 µm) and potato (12-75 µm) (6). Granule size has a major influence on starch end uses. Larger granules are associated with better milling efficiencies and flour yield, improved nutritional properties and specific uses in industries such as the production of paper (7–9).

The size of starch granules is influenced by the timing and location of starch granule initiation. In the Triticeae endosperm, there is a bimodal distribution of starch granules formed by two waves of granule initiation. The first occurs approximately 4-6 days after flowering in the main body of the amyloplast, which produces large, lenticular A-type granules (10). The second round of granule initiation produces B-type granules which are smaller and spherical. B-type granule initiation occurs approximately 10 days after the A-type granules, and at least partially in amyloplast protrusions/stromules (10–12).

However, the genetic control over starch granule size, and factors limiting starch granule growth, are poorly understood. There is relatively little variation in starch granule size among commercial wheat cultivars, with average B-type granule size ranging from 5.8-8.4 µm, and average A-type granule size ranging 17.6-22.5 µm (13). Also, wild relatives, including *Triticum timopheevii*, *T. zhukovskyi, T. urartu* have granule sizes comparable to or smaller than hexaploid wheat (14). Natural variation is therefore unlikely to be a source of alleles for substantially increasing granule size in wheat.

Genetic engineering provides an alternative means to increase granule size, but relies on the identification and testing of suitable gene targets. We recently hypothesised that space within the amyloplast could be a major constraint for starch granule growth. We therefore mutated the plastid division component PARALOG OF ARC6 (PARC6) to increase amyloplast size in wheat (15). However, the consequent increases in granule size were modest, with a 15-22% increase in A-type granule diameter and a 27-44% increase in B-type granule diameter. The modest effect on granule size after increasing amyloplast size was consistent with results in potato (16, 17).

Another strategy is to reduce the number of starch granule initiations, such that there are fewer granules competing for substrates, allowing individual granules to grow larger. We have recently identified components involved in the starch initiation process in wheat (18, 19). B-GRANULE CONTENT 1 (BGC1) is an important target since it influences both A- and B-type granule initiation. A complete knockout of *BGC1* affects both A-type and B-type granule formation (18, 20). It results in supernumerary granule initiation in early grain development, which grow together to form compound granules (where each granule is the result of multiple initiations) (20, 21). Partially reducing *BGC1* expression, in *bgc1-1* mutants, prevents B-type granule formation with no apparent effects on A-type granule morphology (20).

Here, we successfully engineered wheat with substantial increases in starch granule size by simultaneously increasing amyloplast size while reducing granule initiations. Our study demonstrates space and substrate competition as key constraints limiting granule growth in the endosperm, while presenting a novel starch that can be exploited in both food and industrial applications.

## Results

### Isolation of parc6 bgc1 mutants in durum wheat

To study the interaction between starch granule number and amyloplast size, we crossed the *bgc1-1* mutant with the *parc6* mutant of durum wheat (*T. turgidum cv.* Kronos). The *bgc1-1* mutant has reduced function in *BGC1*, by combining a loss of function mutation in *BGC1-A1* (*TRITD4Av1G198830*) with a missense mutation in *BGC1-B1* (*TRITD0Uv1G034540*); and has reduced starch granule number, particularly in the number of B-type granules (20). The *parc6* mutant has premature stop codon mutations in both homeologs of *PARC6* (*TRITD2Av1G286550*, *TRITD2Bv1G255410*) and increased amyloplast size (15). In the F_2_ generation, we isolated lines with homozygous mutations in all homeologs of *PARC6* and *BGC1,* subsequently referred to as *parc6 dbl bgc1*. We also isolated lines with mutations in both homeologs of *BGC1* but only in a single homeolog of *PARC6* (*PARC6-A1*), which provides an intermediate increase in A-type granule size in comparison to the double mutant defective in both homeologs (15). These lines are referred to as *parc6 sgl bgc1.* For comparison, we also isolated a wild-type sibling control, henceforth called WT segregant (WT seg), and mutants defective in both homeologs of each gene - *parc6* and *bgc1*.

The vegetative growth of all mutants in the glasshouse appeared similar to that of the WT. The *parc6 dbl bgc1* plants were slightly shorter at flowering (Fig 1a), but no differences in plant height were observed at grain maturity (Fig S1). Grain yield per plant in the mutants was not different to the WT seg, although some mutants had a significant difference compared to the true Kronos WT, likely due to the effect of background mutations (Fig 1b). There was a small but significant reduction in thousand grain weight in the *parc6 dbl bgc1* mutant compared to the WT segregant, but this was not significant compared to the Kronos WT (Fig 1c). Visually, the grain morphology of *parc6 dbl bgc1* was comparable to the WT controls (Fig 1d), and there was no significant difference in grain size (quantified as the 2D area) (Fig 1e). Overall, grain morphology in plants with mutations in *parc6* and *bgc1* was similar to the WT.

**Figure 1.**
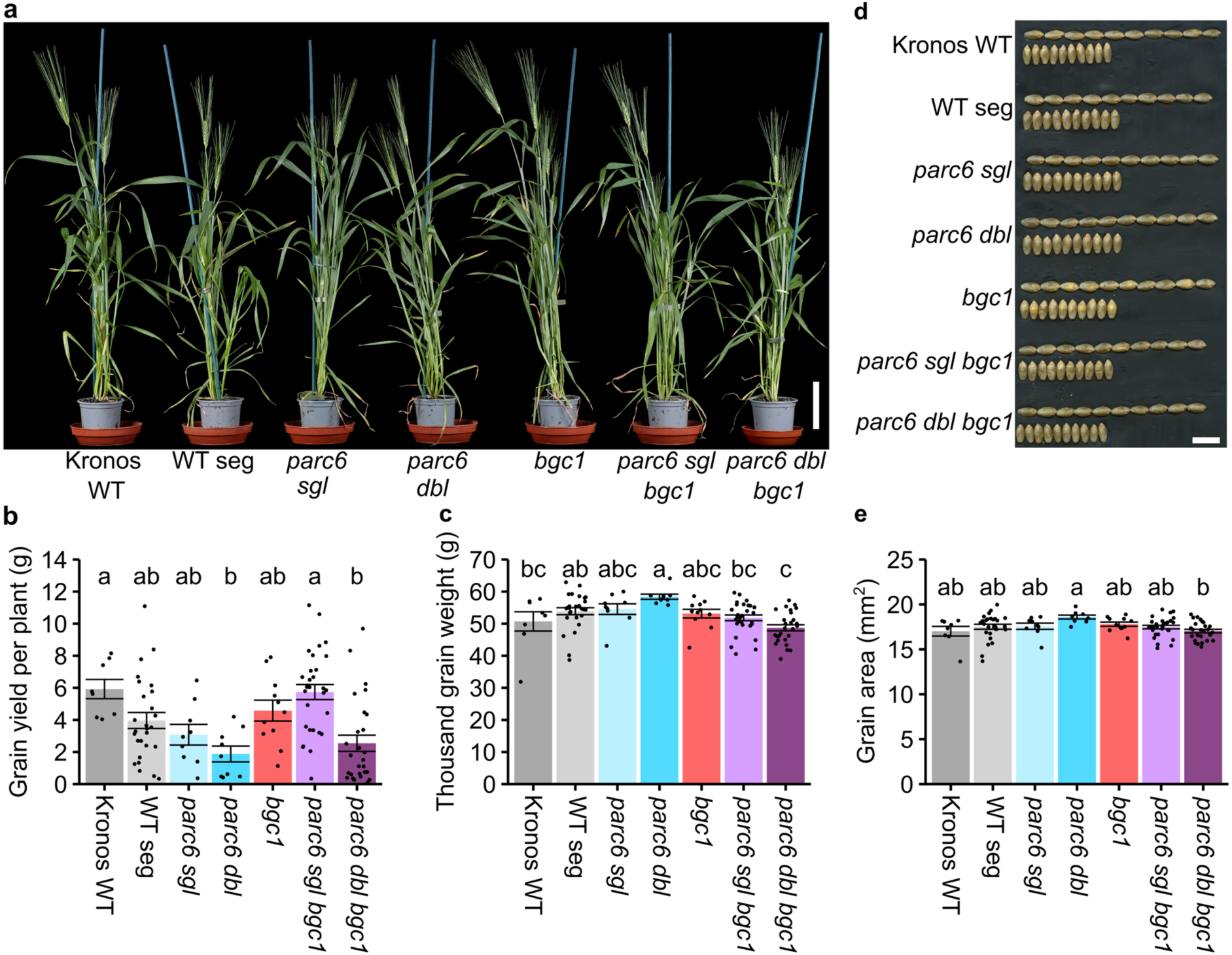
Plant and grain phenotypes of *parc6 dbl bgc1* mutants. (a) Photograph of plants at flowering. Bar = 10 cm. (b) Grain yield per plant. (c) Thousand grain weight. (d) Photographs of grains showing both the dorsal and ventral sides. Bar = 1 cm. (e) Grain size as measured by 2D grain area. In (b), (c) and (e), values are mean ± standard error of the mean, with individual data points shown as black dots. Values with different letters are significantly different under a one-way ANOVA and Tukey’s post hoc test at *P* < 0.05 (*N* = 8-30 per genotype).

### The parc6 dbl bgc1 mutants accumulate massive starch granules

We purified the starch granules from mature grains of the *parc6 bgc1* mutants and examined their morphology. Using scanning electron microscopy (SEM), we saw that *parc6 dbl bgc1* had granules that were extremely large (Fig 2a). The large granules often had abnormal morphology, with creases and ridges on their surfaces rather than a smooth surface. While some smaller granules were present in the mutant, they appeared larger than the B-type granules of the WT. We therefore used a Coulter counter to quantify differences in granule size distribution (Fig 2b). The mutants had altered size distribution curves compared to the bimodal distribution of the Kronos WT and the WT seg. As expected from the SEM, the most extreme shift towards larger size ranges observed in *parc6 dbl bgc1*. All genotypes carrying the *bgc1* mutations had reduced bimodality, with the *bgc1* single mutant having almost no B-type granule peak, and the *parc6 dbl bgc1* having a small shoulder at approximately 10 µm. These are therefore unlikely to represent normal B-type granules as they are much larger than those observed in either the WT segregant or the *parc6* mutants. This shoulder was observed to a lesser extent in *parc6 sgl bgc1*, compared to *parc6 dbl bgc1*.

**Figure 2.**
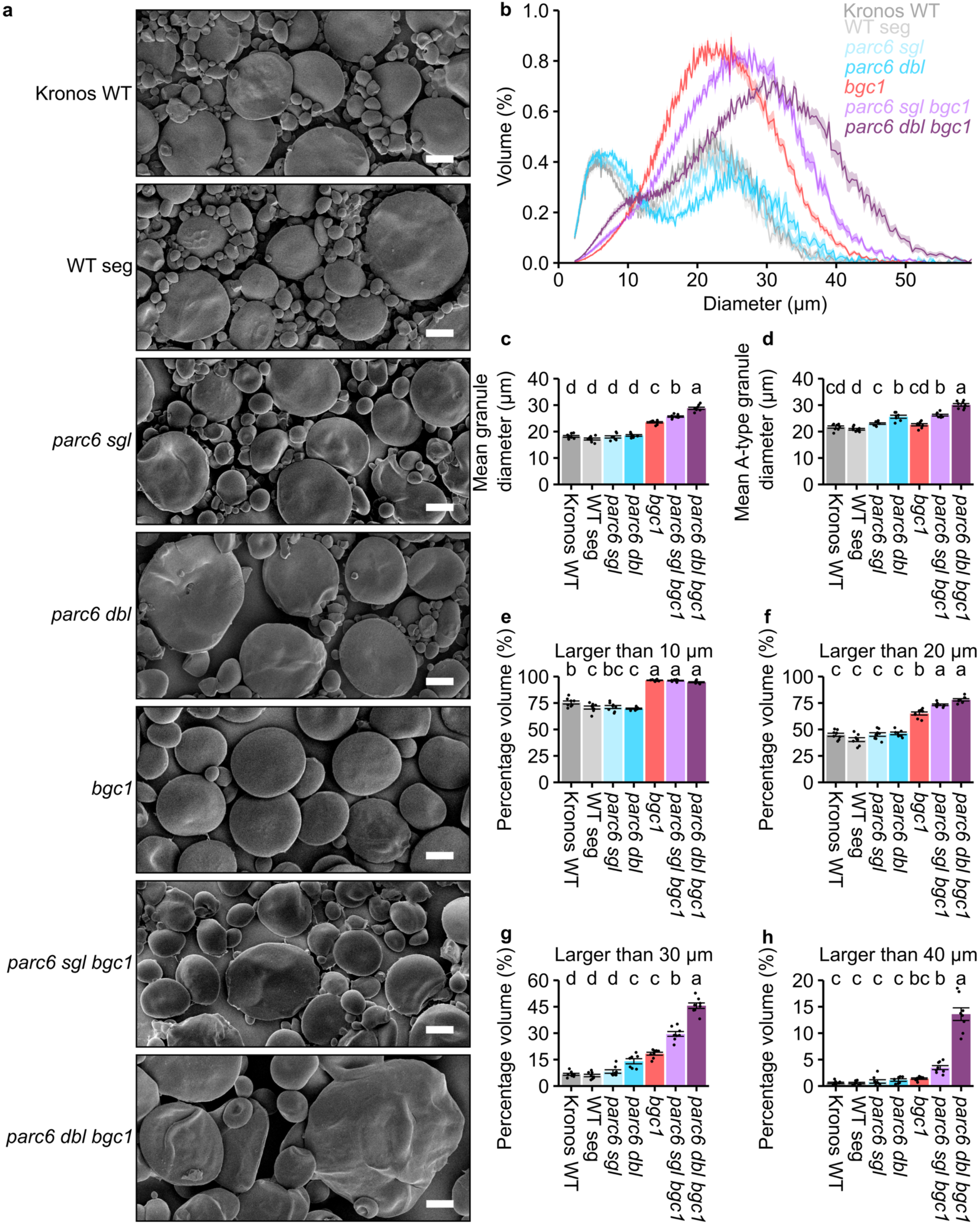
Giant starch granules in *parc6 dbl bgc1* lines. (a) Scanning electron microscopy of purified starch. Bars = 10 µm. (b) Starch was purified from mature grains and a Coulter counter was used to measure size distribution traces. The traces represent mean (solid line) ± standard error of the mean (shading) and have been adjusted for representation on a linear *x* scale. (c) Mean granule diameter, this accounts for both A-type and B-type granules. (d) Mean granule diameter of A-type granules only. (e-h) The percentage (by volume) of granules larger than: (e) 10 µm, (f) 20 µm, (g) 30 µm, (h) 40 µm. Data represent mean ± standard error of the mean, with individual data points shown as black dots, note the altered y-axis scale in (g) and (h) which has been adjusted for clarity. In (c-h) values with different letters are significantly different under a one-way ANOVA and Tukey’s post hoc test at *P* < 0.05 (*N* = 7-8 per genotype).

We then derived granule size data from the size distributions, which demonstrated the striking increases in granule size in the *parc6 bgc1* lines. Given that the genotypes had diverse granule size distributions, we calculated multiple parameters to describe them. Firstly, we calculated the mean granule size (Fig 2c). Among the single gene mutants, the *bgc1* mutation had the strongest effect on increasing average granule size, since removing the small B-type granules greatly increases the average size. However, combining the *parc6* and *bgc1* mutations resulted in further significant increases to the average. We saw a stepwise increase in average granule size from the *parc6 sgl bgc1* to *parc6 dbl bgc1*. Secondly, to quantify the differences in mean A-type granule size, we looked at the diameter at the maximum of the A-type granule peak (Fig 2d). In this aspect, *bgc1* mutation alone had no significant effect on A-type granule diameter. By contrast, *parc6* had a strong, dosage-dependent effect on A-type granule size – with the *parc6 dbl* having a stronger shift than the *parc6 sgl*. However, when combined with *bgc1*, there was an additive effect, where the *parc6 dbl bgc1* had a large shift in A-type granule size that was 44% greater than the WT seg. To further quantify this shift, we calculated the percentage volume of granules above size thresholds (Fig 2e-h). All genotypes containing the *bgc1* mutation have significantly more granules measuring more than 10 µm (Fig 2e), reflecting the reduction in B-type granule formation. The greatest difference among the genotypes was observed in granules greater than 30 µm (Fig 2g). The *parc6 dbl bgc1* mutant had almost half of its total starch volume in granules larger than 30 µm, which was significantly greater than in the WT seg (7%), *parc6 dbl* (14%) and *bgc1* (18%). Even considering granules larger than 40 µm, the *parc6 dbl bgc1* had a substantial amount in this fraction (14%), while all other genotypes had only 0.6-3.5% in this fraction (Fig 2h). All these changes in granule size distribution occurred without any significant effects on starch content, amylopectin chain length structure and amylose content (Fig S2).

Since granule size significantly increased in *parc6 bgc1* mutants in the absence of changes to total starch content, the total number of granules must be decreased. We therefore calculated total granule number using the Coulter counter, and divided this into the number of small (<10 µm) and large (>10 µm) granules (Fig S2). As expected, the *bgc1* mutation has the strongest effect on granule number, which is mainly due to a large reduction in the number of small granules. When combined with the *parc6* mutation, we also observed a significant decrease in the number of large granules compared to the WT seg. We correlated the number of large granules against mean A-type granule diameter and found a significant negative correlation (r = −0.70). This suggests that the number of large granules is decreasing to compensate for the increase in granule size.

We investigated whether the large granule phenotype of *parc6 dbl bgc1* was retained in the field. We grew three replicated 1 m^2^ plots for WT seg, *parc6 sgl*, *parc6 dbl*, *bgc1*, *parc6 sgl bgc1* and *parc6 dbl bgc1*. The phenotypes of these plants were comparable to those grown under glasshouse conditions (Figs S3-5). Surprisingly, the yield of the WT seg was the lowest and, with the exception of the *parc6 dbl*, all mutants yielded significantly more grains than the WT seg (Fig S3a). In contrast to the glasshouse grown grains, there was no significant difference in thousand grain weight between the *parc6 dbl bgc1* and WT seg plants (Fig 1c, Fig S3b). This suggests that the small decrease in thousand grain weight observed in *parc6 dbl bgc1* plants grown in the glasshouse may not be reproduced under field conditions.

Taken together, combining *parc6* and *bgc1* mutations generated a range of novel genotypes carrying substantial variation in granule size distributions, including the giant granules in *parc6 bgc1* that were substantially larger than those observed in either mutant alone.

### Larger amyloplasts and reduced granule initiations facilitate giant granules in parc6 dbl bgc1

We hypothesised that in *parc6 bgc1* mutants, the large amyloplasts would have a reduced number of granules relative to the *parc6* mutants, hence each individual granule would have more space and substrates to grow. We previously reported that amyloplasts in *parc6* mutants contain multiple large granules (15), which we confirmed here using transmission electron microscopy (TEM) at 15 days post flowering (DAF) (Fig 3). However, we did not observe multiple large granules in amyloplasts from *parc6 dbl bgc1* lines, and the starch granules were strikingly larger than those in the WT seg and the *parc6* and *bgc1* single mutants, consistent with our Coulter counter data (Fig 2b). In the *parc6 dbl*, the amyloplast envelope was often not closely associated with the developing granule, suggesting availability of stromal space, as previously observed (15). However, in *parc6 dbl bgc1*, the amyloplast membrane appeared more tightly associated with the growing granule.

**Figure 3.**
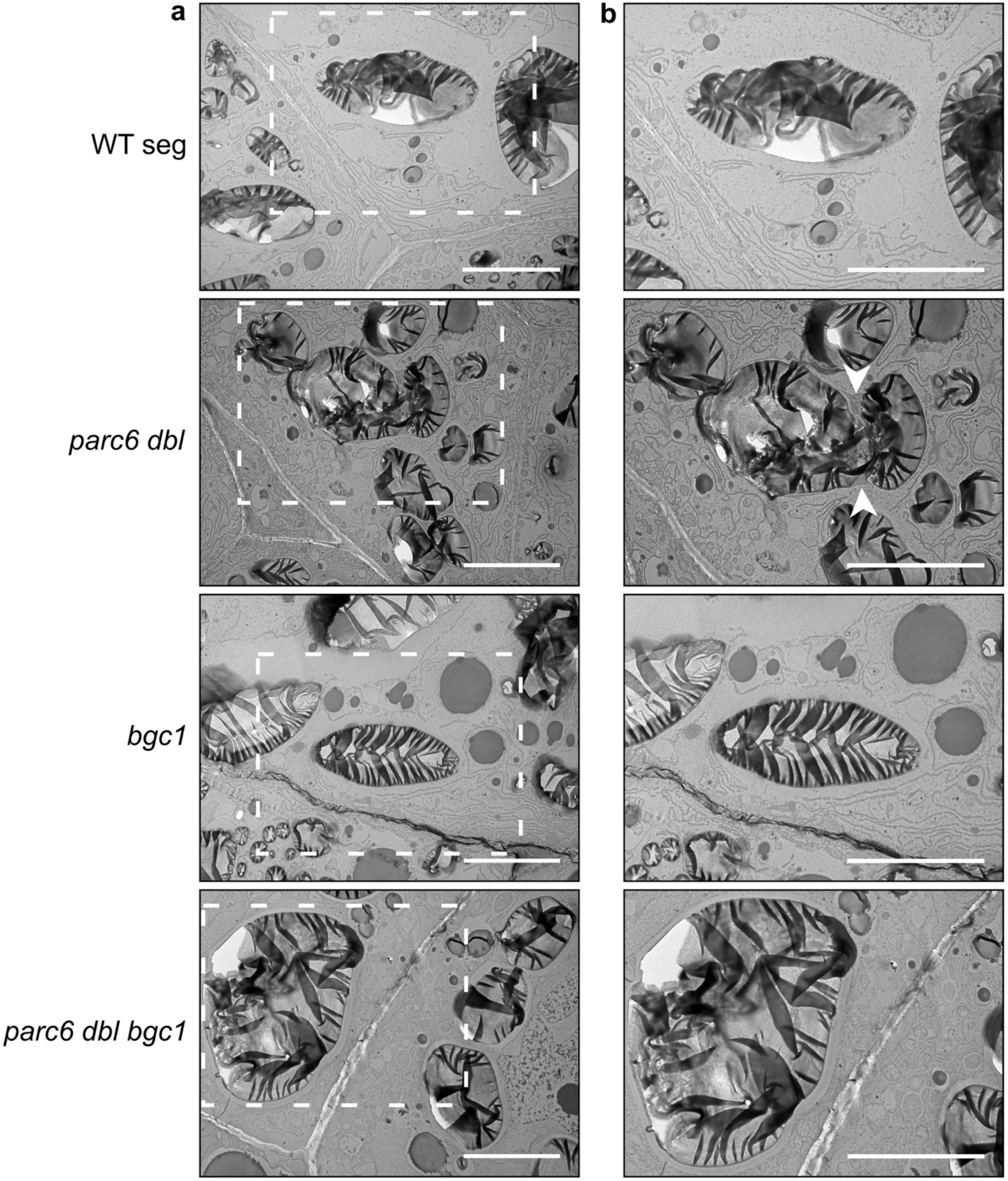
Starch granules in amyloplasts of *parc6 bgc1* mutants. (a-b) Transmission electron microscopy (TEM) images of endosperm sections of developing grain at 15 days after flowering (DAF). Bars = 10 µm. White dashed boxes in (a) represent the area which has been enlarged in (b). Where visible, the periphery of amyloplasts is indicated with white arrows.

### Purified starch from parc6 bgc1 mutants have altered physiochemical properties linked to differences in A-type granule size

We previously demonstrated that A-type granule size has a major influence on physiochemical properties of starch (22). We therefore investigated whether the granule size increases in our wheat lines altered the pasting properties of starch using a Rapid Visco Analyser (RVA). Lines carrying *bgc1* mutations consistently had greater peak viscosities and lower pasting temperatures compared to the WT seg (Fig S6). The *parc6 bgc1* starch did not have statistically significant differences in rheological properties compared to *bgc1* starch alone, likely due to the low statistical power in the experiment. However, multiple rheological properties significantly correlated with mean A-type granule size (Fig 4). The strongest relationship was the negative correlation between A-type granule size and pasting temperature (r = −0.87, *P* < 0.001). A-type granule size also positively correlated with peak viscosity (r = 0.80, *P* < 0.001), breakdown viscosity (r = 0.76, *P* < 0.001) and final viscosity (r = 0.74, *P* < 0.001). We also measured thermal properties using a Differential Scanning Calorimeter (DSC) (Fig S7). Consistent with the RVA, the largest effect was observed with *bgc1* mutations, which reduced gelatinisation temperature by ∼2°C. However, unlike the RVA, the gelatinisation temperature measured by DSC did not correlate well to A-type granule size. Also, none of the different genotypes significantly differed in starch swelling power (Fig S8).

**Figure 4.**
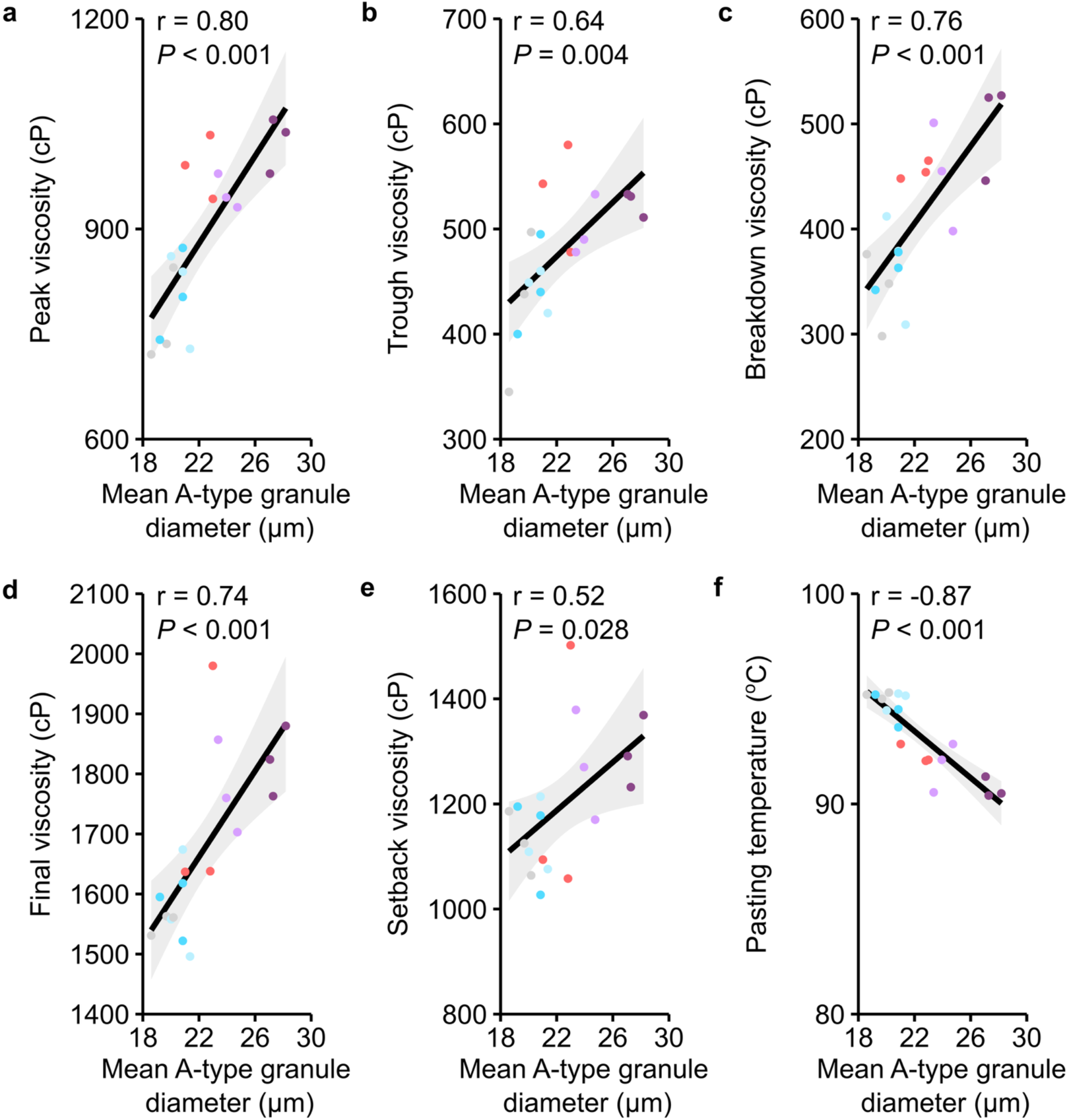
Large starch granules from *parc6 bgc1* mutants have altered pasting temperatures and viscosities. Purified starch (2 g in 25 mL water) was cooked and cooled on the Rapid Visco Analyser (RVA) and parameters from the viscographs were obtained and correlated against mean A-type granule diameter. (a) Peak viscosity, (b) trough viscosity, (c) breakdown viscosity, (d) final viscosity, (e) setback viscosity, (f) pasting temperature. The black line represents a linear model between the parameters with the grey shading representing 95% confidence intervals. A Pearson correlation test was conducted and the correlation coefficients (r), and corresponding *P* value are on the top left of each panel.

Since larger granules were previously associated with reduced digestion rates *in vitro* (23, 24), we also tested starch digestion properties in the *parc6 bgc1* lines. However, we did not observe any differences between genotypes in resistant starch content, or in the digestion kinetics of native starch granules with α-amylase *in vitro* (Fig S9). Therefore, the range of granule sizes we examined does not seem to influence digestion properties under these assay conditions. However, it remains to be determined whether there are differences in digestibility in a cooked food product or in different food matrices.

## Discussion

### Engineering of giant starch granules in wheat

Our work demonstrates two interlinked factors limiting starch granule size: the space available for starch granule growth (amyloplast size), and the number of granules competing for the space and substrates (the number of initiated granules). We found that to engineer substantial increases in granule size, both parameters had to be targeted through mutations in independent gene targets. The necessity of two gene mutations explains why such giant starch granules were never achieved through traditional mutant screening and plant breeding.

We previously noted that the large A-type granules in modern wheat varieties have an average size that ranges 17.6-22.5 µm (13). However, since A-type starch granules follow normal or log-normal distributions, granules at the maximum end of the distribution can reach up to 30 µm, but they are extremely rare. In the WT, only 6% of starch were in granules larger than 30 µm, and almost none above 40 µm (Fig 2). By contrast, our *parc6 dbl bgc1* line that had the most extreme increase in granule size had almost half of its starch volume in granules larger than 30 µm, and more than 14% in granules larger than 40 µm (Fig 2). Such a large shift in granule size distribution is highly novel, and has never been observed in the Triticeae (14, 25), nor in other cereals.

We suggest that the large granules form in *parc6 dbl bgc1* because they have space to grow in the enlarged amyloplasts, without competing with other granules within the same amyloplast for space and substrates. Our previous work demonstrated modest increases in A-type and B-type granule size could be achieved by increasing amyloplast size by mutating *PARC6* (15). However, increasing amyloplast size also increased the number of granules per amyloplast, with many containing multiple A-type granules (15) (Fig. 3). This parallels observations in Arabidopsis plastid division mutants, where increasing chloroplast size also increased the number of granules contained within each chloroplast, such that the number of granules per stromal area remains constant (26). Thus, in Arabidopsis, increasing chloroplast size did not affect average granule size (27). Notably, *parc6* mutants of wheat also have increased production of B-type granules (15). Having multiple A-type granules and B-type granules within each amyloplast competing for space and substrates would substantially limit granule growth, explaining why *parc6* mutants do not have more substantial increases in granule size. The size increase achieved by combining *parc6* with the *bgc1* mutation, which greatly reduced the number of initiations of both A- and B-type granules (Fig S2), is consistent with this hypothesis (Figs 2, S4, S5). Interestingly, preventing B-type granule initiation alone (i.e., in the *bgc1* mutant alone) does not lead to detectable increases in A-type granule size (20) (Fig 2), despite no apparent changes in total starch content (28)(Fig S2). Therefore, redirecting substrates from B-type granule growth does not seem to be sufficient for creating detectable increases in A-type granule size.

Despite the presence of very large granules in the *parc6 bgc1* lines, the size distribution curves are broad with many granules smaller than 20 µm in size (Fig S4). Surprisingly, there was also a shoulder around 10 µm in the size distribution trace of *parc6 dbl bgc1*, pointing to a population of smaller granules – although these are much larger than normal B-type granules. This was unexpected as the *bgc1-1* mutation was thought to almost completely suppress B-type granule initiation (20). However, the *bgc1-1* mutation is a hypomorphic mutant, because it combines a knockout mutation in *BGC1-A1* with a missense mutation in *BGC1-B1*. It was important to use this hypomorphic mutation in our study, since the full knockout mutant in *BGC1* has defective granule morphology due to compound granule formation. However, a consequence of this is that *bgc1-1* may not completely suppress B-type granule formation. Residual B-type granule formation may not be detectable in *bgc1-1*, but may become detectable in a *parc6* background where B-type granule formation is elevated. The reason why B-type granules are more frequent in *parc6* is currently cannot be explained, but suggests an influence of amyloplast size in the process.

We also do not yet understand why the large granules in *parc6 dbl bgc1* lines adopt abnormal morphologies. It is unlikely to be purely related to their size - starch granules from root/tuber crops like potatoes and yams naturally grow much larger (up to 100 µm) without morphological defects (6). In *parc6* mutants, these abnormal morphologies occur from early grain development, when the granules are not likely to be space limited (15). A-type granules undergo a defined morphogenesis program, where they are initially spherical, and gradually adopt their defined shape (29). It is possible that the enlarged amyloplasts in *parc6* prevents this morphogenesis from occurring, suggesting an important role of amyloplast structure for correct morphogenesis.

### Applications of giant wheat starch granules

Starch granule size is one of the key factors determining starch applications, alongside amylose content and polymer structure. Aside from its major application in food, wheat starch also has industrial uses in the production of paper and corrugated board, cosmetics, biodegradable plastics, and for drilling oil (9). Our approach has created a cereal starch in a size range that was previously not available, naturally or artificially, and the novelty of the material provides an opportunity to find beneficial applications. We propose several benefits of increasing granule size in the Triticeae. Firstly, larger starch granules are associated with better milling efficiencies and flour yields (7). For the food industry, increasing the amount of larger starch granules reduces overall B-type granule content (the total starch volume composed of B-type granules) which is favourable for producing high-quality bread and noodles (30–32). In many industries, wheat starch is often separated into small granule and large granule fractions with a hydrocyclone (9). The large granule fraction is useful in the production of materials such as carbonless copy paper (33).

Crucially for applications, the granule size phenotypes of *parc6 bgc1* lines were highly reproducible in field or glasshouse conditions, without consistent differences in grain yield per plant or per plot (Fig S3). This is in contrast to the rice *parc6* mutant, which has reduced grain yield in field conditions (34). These data show promise of *parc6 bgc1* lines being grown in the field at scale and integrated into wheat breeding programs, although proper field trials in different environments will be needed.

We also examined the rheological properties of the starch granules. In both RVA and DSC, the gelatinisation peak shifted to lower temperatures in starch containing the *bgc1* mutation (Figs S6, S7). This is unsurprising given that B-type granules are generally reported as having a higher peak gelatinisation (35–38). Interestingly, there was no correlation between granule size and peak temperature in DSC, suggesting there are structural differences in A-type and B-type granules which are independent of granule size. By contrast, in RVA, A-type granule size significantly correlated with pasting and viscosity parameters (Fig 4). A key distinction between RVA vs. DSC is that RVA results are influenced not only by the crystalline melt temperature, but also by granule swelling properties and particle packing. This is in agreement with the results of Fahy, Chen and Seung (22) who suggested that A-type granule size is the major factor influencing starch granule pasting.

Larger granules digest more slowly in vitro as they have a smaller surface area to volume ratio (39, 40). Despite the significanct increase in granule size, we saw no differences in granule digestibility or resistant starch content (Fig S9). This could be due to the morphological defects in the large granules of *parc6 dbl bgc1*, including ridges on the granule surface that would increase surface area for enzyme binding (Fig 2a). However, it is important to note that the digestion of raw wheat starch granules is not relevant to nutrition, as starch is consumed after cooking in a food matrix. Many starch granules remain intact after cooking, for example in pasta where granules are surrounded by the gluten network (41). Thus, it is important to now assess the effect of increased granule size in our wheat lines on nutritional properties in a cooked food product.

## Materials and Methods

### Plant material and growth conditions

The *parc6* mutant has been previously described in Esch*, et al.* (15). In this study, double homeolog mutants are referred to as *parc6 dbl* (*aabb*), and the A homeolog single mutant is referred to as *parc6 sgl* (*aa*BB). For simplicity, only the A homeolog single mutant was included in the analysis as Esch*, et al.* (15) found no differences between the A homeolog and B homeolog single mutants. The *bgc1-1* mutant was previously described in Chia*, et al.* (20) – for simplicity this has been shortened to *bgc1* in the text and figures. The *parc6 dbl* was crossed to *bgc1* plants and in subsequent generations, we isolated double homeolog mutants for either *parc6* (*parc dbl*), or bgc1 (*bgc1*), or double homeolog mutants for both genes (*parc6 dbl bgc1*). Single homeolog mutants for *parc6* (*parc6 sgl*, *aa*BB) and single homeolog mutants for *parc6* and double homeolog mutants for *bgc1* (*parc6 sgl bgc1*) were also isolated. Corresponding wild type segregants (WT seg) were also isolated as controls. Plants were genotyped using KASP genotyping using primers described in Chia*, et al.* (20) and Esch*, et al.* (15). In all experiments, a wild type *T. turgidum cv*. Kronos (Kronos WT) was also included.

Plants grown in the glasshouse received a minimum of 16 hours light at 20°C and 16°C during the dark. Plants were grown in 9 cm pots filled with John Innes cereal mix (65% peat, 25% loam, 10% grit, 3 kg/m^3^ dolomitic limestone, 1.3 kg/m^3^ PG mix, 3 kg/m^3^ osmocote extract).

For bulking grains for starch physicochemical analyses, we grew plants in the field during spring/summer 2025 at the John Innes Centre Dorothea de Winton Field Station (Church Farm, Norfolk, UK, 52°37′49.2″ N 1°10′40.2″ E). Plants were grown with standard agronomic practices in 1 m^2^ plots with 3 plots per line.

### Grain and plant morphometrics

Grain number and grain size traits were quantified with the MARViN seed analyser (Marvitech GmbH, Wittenburg). For field grown plants, only a fraction of harvested grains (1245-1582 individual grains per plot) were analysed. For glasshouse-grown plants, all harvested grains were examined. Grain yield was quantified as the total weight of grains harvested: For glasshouse-grown samples this was measured using the MARViN, and for field-grown samples total grain weight was measured with a fishing scale (Meilen). Plant height and tiller number was measured in mature plants before grains were harvested.

### Starch purification, granule morphology and size distribution

For each starch purification, three mature grains were soaked overnight in ddH_2_O at 4°C and homogenised with a mortar and pestle with excess ddH_2_O. The homogenates were filtered through a 100 µm filter (pluriStrainer, pluriSelect) and centrifuged at 3,000*g* for 5 mins. The pellet was resuspended in water and centrifuged at 2,500*g* for 5 min on a cushion of 90% (v/v) Percoll, 50 mM Tris-HCl pH 8.0. The pellet was washed twice with 50 mM Tris-HCl, pH 6.8, 10 mM EDTA, 4% SDS (v/v), and 10 mM DTT and a further three times with ddH_2_O, before the purified starch was resuspended in ddH_2_O. To produce sufficient starch from field-grown grains for analysis on the Rapid Visco Analyser and α-amylase digest assays, this method was scaled up. A cyclone mill (Retsch) with a 2 mm sieve and a speed of 14,000 rpm was used to produce wholemeal flour, which was resuspended in excess ddH_2_O. The homogenates were filtered through muslin and miracloth and centrifuged at 3,124*g* for 5 min. The pellet was spun through a Percoll gradient and washed in the same way as described above.

Granule morphology was examined via scanning electron microscopy. Diluted purified starch was mounted on the surface of an aluminium pin stub using double-sided adhesive carbon discs (Agar Scientific Ltd, Stansted, Essex). The stubs were sputter coated with approximately 8 nm gold in a high-resolution sputter coater (Agar Scientific Ltd) and transferred to a FEI Nova NanoSEM 450 (FEI, Eindhoven, The Netherlands). The samples were viewed at 3 kV and digital TIFF files were stored.

For analysis of granule size, purified starch was resuspended in Isoton II electrolyte solution (Beckman Coulter, Indianapolis) and particle size distributions were measured using a Multisizer 4e Coulter counter (Beckman Coulter) with a 100 µm aperture tube and counting a minimum of 100,000 particles per sample. All Coulter counter traces shown here have been transformed to account for logarithmic bin sizing during data collection and are represented on a linear scale. To calculate mean A-type granule diameter, a rolling average with a sliding window of 50 was applied to the data, and the maxima of the peak > 10 µm was determined.

### Starch and amylose quantification

For quantification of total starch content, two mature grains were ground to wholemeal flour using a Geno/Grinder (SPEX CertiPrep™) at 1500 rpm for 10 min. Flour (5-10 mg) was dispersed in 20 μL of 80% (*v/v*) ethanol and incubated with 500 μL of thermostable α-amylase in 100 mM sodium acetate buffer (pH 5.0) at 80°C for 20 min with regular shaking of the samples to digest the starch into maltodextrins. Amyloglucosidase was added and samples were incubated at 5°C for 35 min to digest the maltodextrins into glucose. The samples were centrifuged at 3220*g* for 10 min. The supernatant (5 μL) was used in a spectrophotometric hexokinase/glucose-6-phosphate dehydrogenase assay to measure glucose. All the enzymes and reagents for these steps were from the Total Starch Assay Kit (K-TSHK, Megazyme).

For quantification of amylose, 1 mg of purified starch was dissolved in 200 μL water and 200 μL of 2 M NaOH and left to incubate at room temperature overnight. The solution was neutralised to pH 7 with 1 M HCl. The solution (5 μL) was diluted in 220 μL of water and 25 μL Lugol’s iodine solution (Sigma Life Science) and incubated at room temperature for 10 min. The absorbance at 535 nm and 620 nm and the apparent amylose content was estimated as outlined by Washington, Box, Karakousis and Barr (42).

### Electron microscopy analysis of developing grains

Developing grains (15 days after flowering) were harvested into 2.5% (w/v) glutaraldehyde in 0.05 M sodium cacodylate, pH 7.4. The grains were post-fixed with osmium and dehydrated with an ethanol series. Grains were embedded in LR white resin with an EM TP embedding machine (Leica). Ultrathin sections (∼90 nm) were prepared using a diamond knife and a Leica UC7 ultramicrotome. Sections were transferred to formvar and carbon coated 200 mesh copper grids and stained with 2% (w/v) uranyl acetate for 1 hour followed by 1% (w/v) lead citrate for 1 min. Grids were washed with distilled water and airdried and imaged on a FEI Talos 200C transmission electron microscope (FEI UK Ltd, Cambridge, UK) at 200 kV and imaged using a Gatan OneView 4 K × 4 K digital camera (Gatan, Cambridge, UK) to record DM4 files.

### Rheological, Thermal and Swelling analyses

Rapid Visco Analysis (RVA) was carried out on an RVA Tecmaster instrument (Perten, 174 Waltham, MA, USA) with 2 g purified starch in 25 mL of water. The preinstalled general pasting method was used according to AACC 175 Method 76-21.

Differential Scanning Calorimetry (DSC) analysis was carried out with a multi-cell DSC (TA Instruments). Purified starch (100 mg) was weighed into Hastelloy ampoules. Samples were heated from 10°C to 150°C in a furnace purged by Nitrogen gas at a flow rate of 50 mL per min at a scan rate of 0.5 °C per min with a pre-equilibration step of 600 seconds.

Swelling power was determined based on the method of Howard*, et al.* (43). Briefly, purified starch (10 mg) was suspended in 1 mL of water and heated at 80°C for 20 min in a thermomixer set to 750 rpm. Samples were cooled at 22°C for 5 min, and spun at 1500 x *g* at 22°C for 5 min. The supernatant was removed and the weight of the starch pellet measured. Swelling power is reported as the fractional increase in the weight of the starch.

### Resistant starch content and digestibility

For determination of resistant starch, the K-RSTAR kit (Megazyme) was used. The method was scaled down to use as starting material 10 mg of flour, produced using a cyclone mill (Retsch) with a 1 mm sieve, and all assay volumes were reduced by 10X.

The α-amylase digestion assays were conducted with 100 mg of purified starch in 10 mL of phosphate buffered saline, samples were rotated at 37°C for 30 min before the addition of 4 U of porcine pancreatic α-amylase (Sigma-Aldrich, EC 3.2.1.1, A6255). Aliquots (100 µL) were taken at set time points and the reaction stopped with 100 µL of 0.3 M sodium carbonate. The concentration of released maltose was determined with the *p*-hydroxybenzoic acid hydrazide (PAHBAH, CAS No.5351-23-5) (44)

### Statistics and data visualisation

All statistical analyses were conducted with RStudio (version 2023.12.1, build 402). The statistical tests used, and any post hoc tests, are specified in the legend or text. Parametric tests were used, except in cases where the data did not follow a normal distribution, as determined by Shapiro-Wilk tests and visual inspection of quantile-quantile plots. In these cases, non-parametric tests were used instead. The ‘ggplot2’ package in RStudio was used to generate all graphs.

## Author Contributions

DS conceived the study. RM, LE, FW and DS designed the study. RM, LE, QYN, KP, RS and BF conducted the experiments and analysed data. RM and DS wrote the paper with input from all authors.

## Acknowledgements

We thank JIC Horticultural Services for providing growth facilities and maintenance of plant material, JIC Bioimaging for providing access to microscopes, and the JIC Genotyping service for conducting KASP genotyping. The *bgc1-1* mutant was originally provided by Tansy Chia (NIAB) and Kay Trafford (JIC). This work was funded through a John Innes Foundation (JIF) Chris J. Leaver Fellowship (to D.S), Leverhulme Trust Research Project grant RPG-2019-095 (to D.S), a Genetics Society Summer Studentship (to K.P), and Biotechnology and Biological Sciences Research Council (BBSRC, UK) research grants BB/W015935/1, BB/W01632X/2, BB/Z517501/1 (all to D.S) and UKRI1923 (to R.M and D.S), BBSRC Biofortification Hub grant BB/X010864/1-012 (to D.S), and BBSRC Institute Strategic Programme grants BB/X01097X/1 and BB/X011003/1 (to the John Innes Centre). The John Innes Centre Bioimaging Platform is supported by BBSRC grant BB/CCG2240/1.

## Supplemental Materials

**Figure S1.**
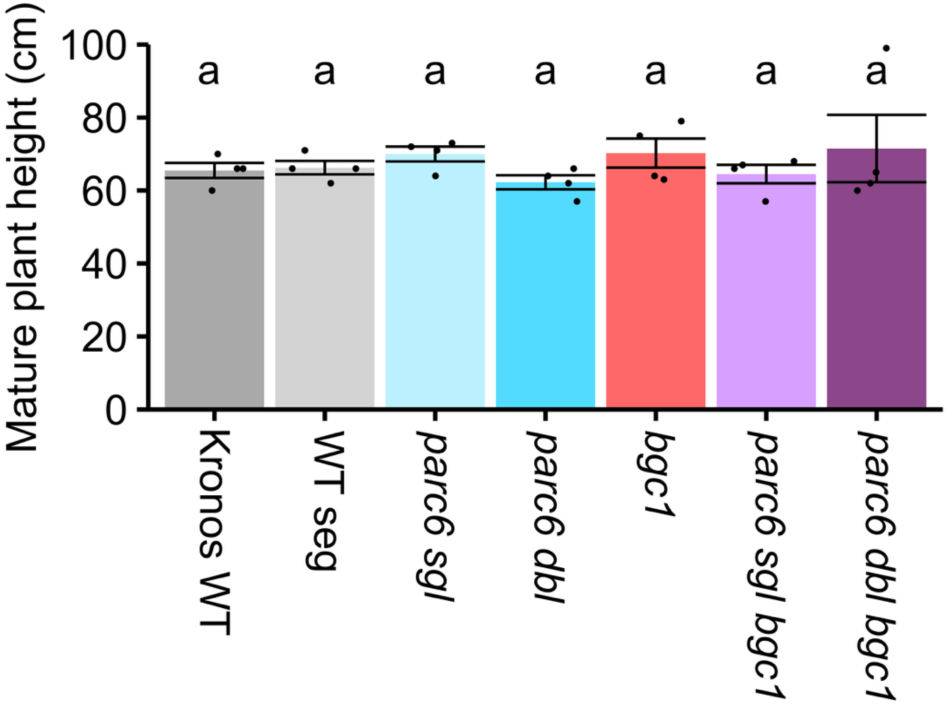
There is no difference in the height of mature *parc6 bgc1* mutants. Data are presented as means ± standard error of the mean, with individual data points shown as black dots. Values with different letters are significantly different under a one-way ANOVA and Tukey’s post hoc test at *P* < 0.05 (*N* = 4 per genotype).

**Figure S2.**
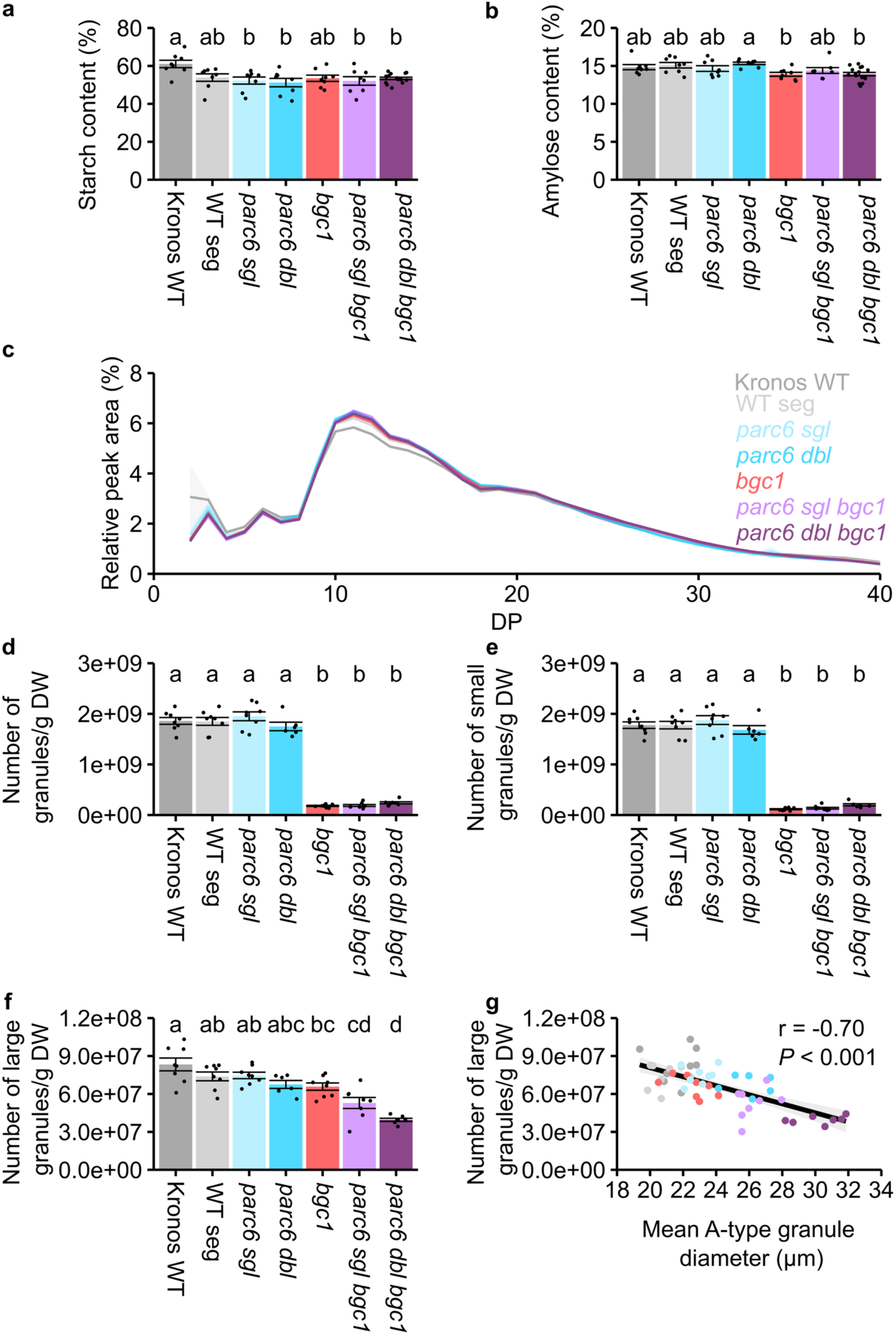
Amylose content, total starch content and granule number in *parc6 bgc1* mutants. (a) Total starch content of wholewheat flour. (b) Amylose content of purified starch. Data in (a)-(b) are presented as means ± standard error of the mean, with individual data points shown as black dots. Values with different letters are significantly different under a one-way ANOVA and Tukey’s post hoc test at *P* < 0.05 (*N* = 8 per genotype). (c) Chain length distribution of purified starch, data are represented as means (solid lines) ± standard error of the mean (shading), (*N* = 3-4 per genotype). (d) Starch granule number in mature grains. Starch was purified, and the number of granules was determined using a Coulter counter running in volumetric mode. Values are expressed relative to the dry weight of the grain. (e) Starch granule number from (d) but only granules <10 µm were counted. (f) Starch granule number from (d) but only granules >10 µm were counted. In (d-f), data are presented as means ± standard error of the mean, with individual data points shown as black dots. Values with different letters are significantly different under a one-way ANOVA and Tukey’s post hoc test at *P* < 0.05 (*N* = 6-8 per genotype). (g) The number of large granules (f) correlated against mean A-type granule diameter (from Fig 2d). The black line represents a linear model between the parameters with the grey shading representing 95% confidence intervals. A Pearson correlation test was conducted and the correlation coefficient (r) and *P* value are on the top right of (g).

**Figure S3.**
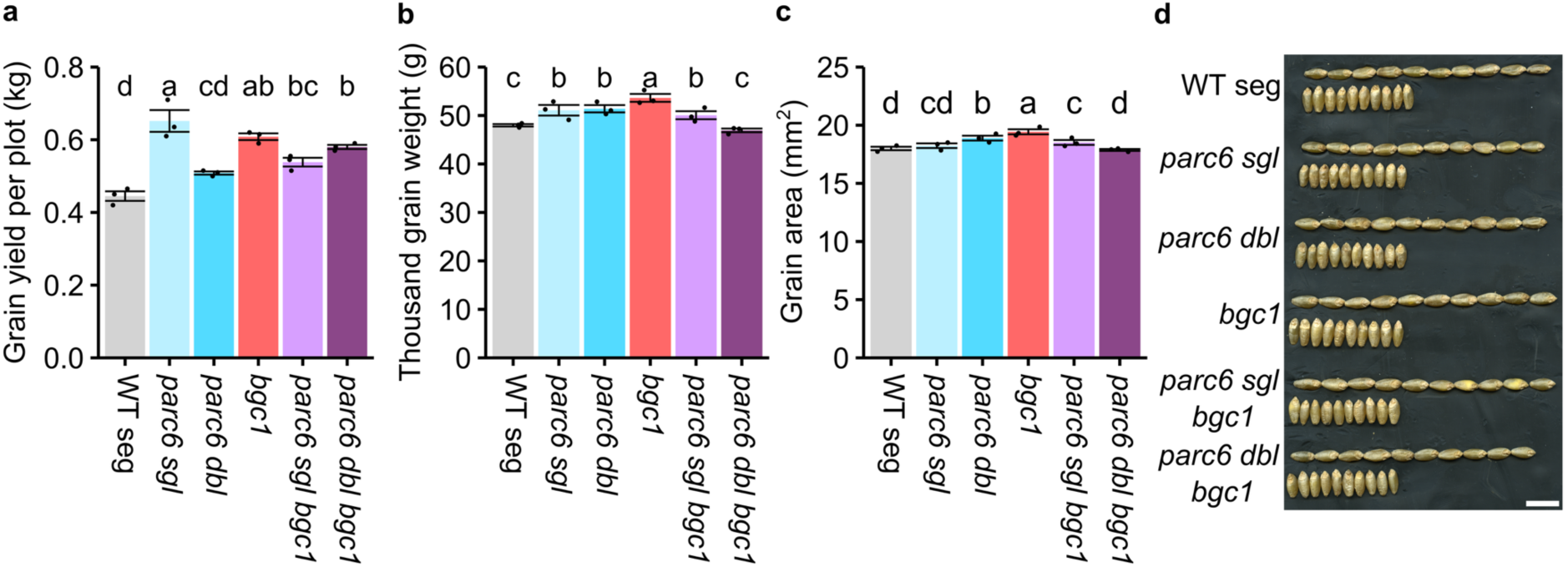
Grain phenotypes of field-grown *parc6 bgc1* mutants. (a) Grain yield per 1m^2^ plot. (b) Thousand grain weight. (c) Grain size as measured by 2D grain area. (d) Photographs of grains showing both the dorsal and ventral sides. Bar = 1 cm. (e) In (a), (b) and (c) data are presented as means ± standard error of the mean, with individual data points shown as black dots. Values with different letters are significantly different under a one-way ANOVA and Tukey’s post hoc test at *P* < 0.05 (*N* = 3 per genotype).

**Fig S4.**
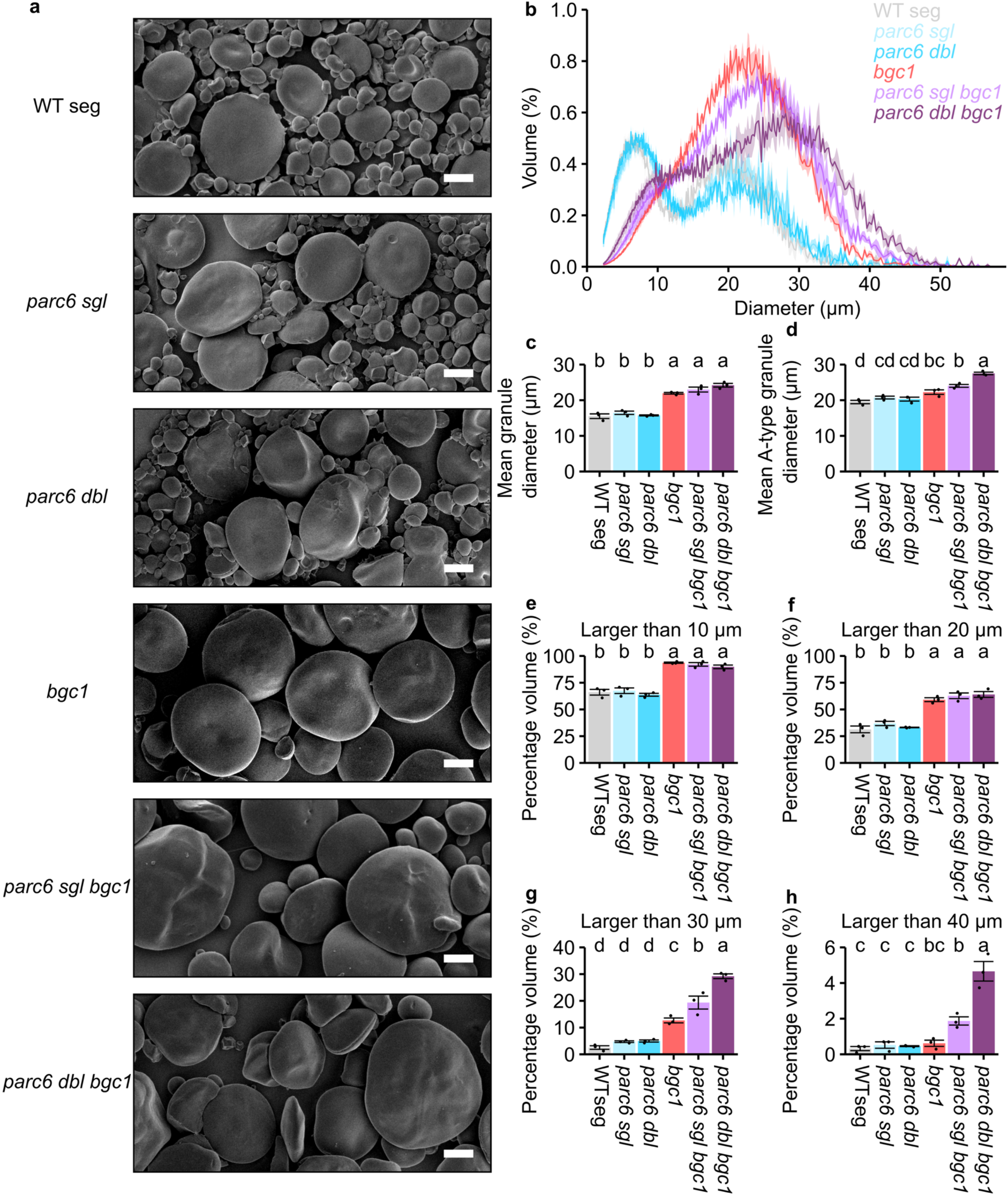
Granule size phenotypes are reproducible in field-grown *parc6 bgc1* mutants. (a) Scanning electron microscopy of purified starch. Bar = 10 µm. (b) Starch was purified from mature grains and a Coulter counter was used to measure size distribution traces. The traces are displayed as means (solid lines) ± standard error of the mean (shading) and have been adjusted for representation on a linear *x* scale. (c) Mean granule diameter, this accounts for both A-type and B-type granules. (d) Mean granule diameter of A-type granules only. (e-h) The percentage (by volume) of granules larger than: (e) 10 µm, (f) 20 µm, (g) 30 µm, (h) 40 µm. Data are presented as means ± standard error of the mean, with individual data points shown as black dots, note the altered y-axis scale in (g) and (h) which has been adjusted for clarity. Values with different letters are significantly different under a one-way ANOVA and Tukey’s post hoc test at *P* < 0.05 (*N* = 3 per genotype).

**Fig S5.**
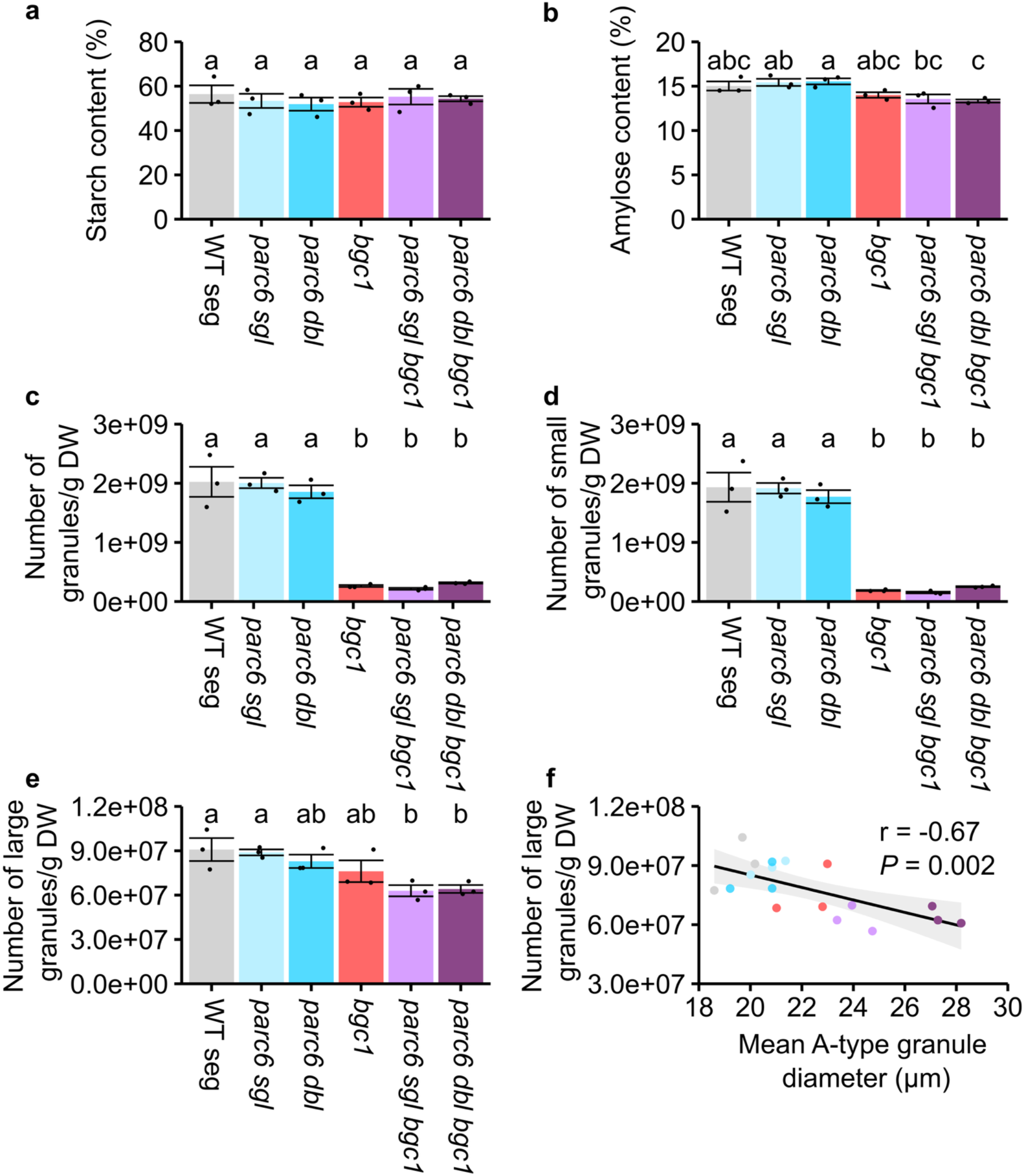
Amylose content, total starch content and granule number in field-grown *parc6 bgc1* mutants. (a) Amylose content of purified starch. (b) Total starch content of wholewheat flour. Values with different letters are significantly different under a one-way ANOVA and Tukey’s post hoc test at *P* < 0.05 (*N* = 3 per genotype). (c) Starch granule number in mature grains. Starch was purified, and the number of granules was determined using a Coulter counter running in volumetric mode. Values are expressed relative to the dry weight of the grain. (d) Starch granule number from (c) but only granules <10 µm were counted. (e) Starch granule number from (c) but only granules >10 µm were counted. In (a-e), data are presented as means ± standard error of the mean, with individual data points shown as black dots. Values with different letters are significantly different under a one-way ANOVA and Tukey’s post hoc test at *P* < 0.05 (*N* = 3 per genotype). (f) The number of large granules (e) correlated against mean A-type granule diameter (from Fig S4d). The black line represents a linear model between the parameters with the grey shading representing 95% confidence intervals. A Pearson correlation test was conducted and the correlation coefficient (r) and *P* value are shown.

**Fig S6.**
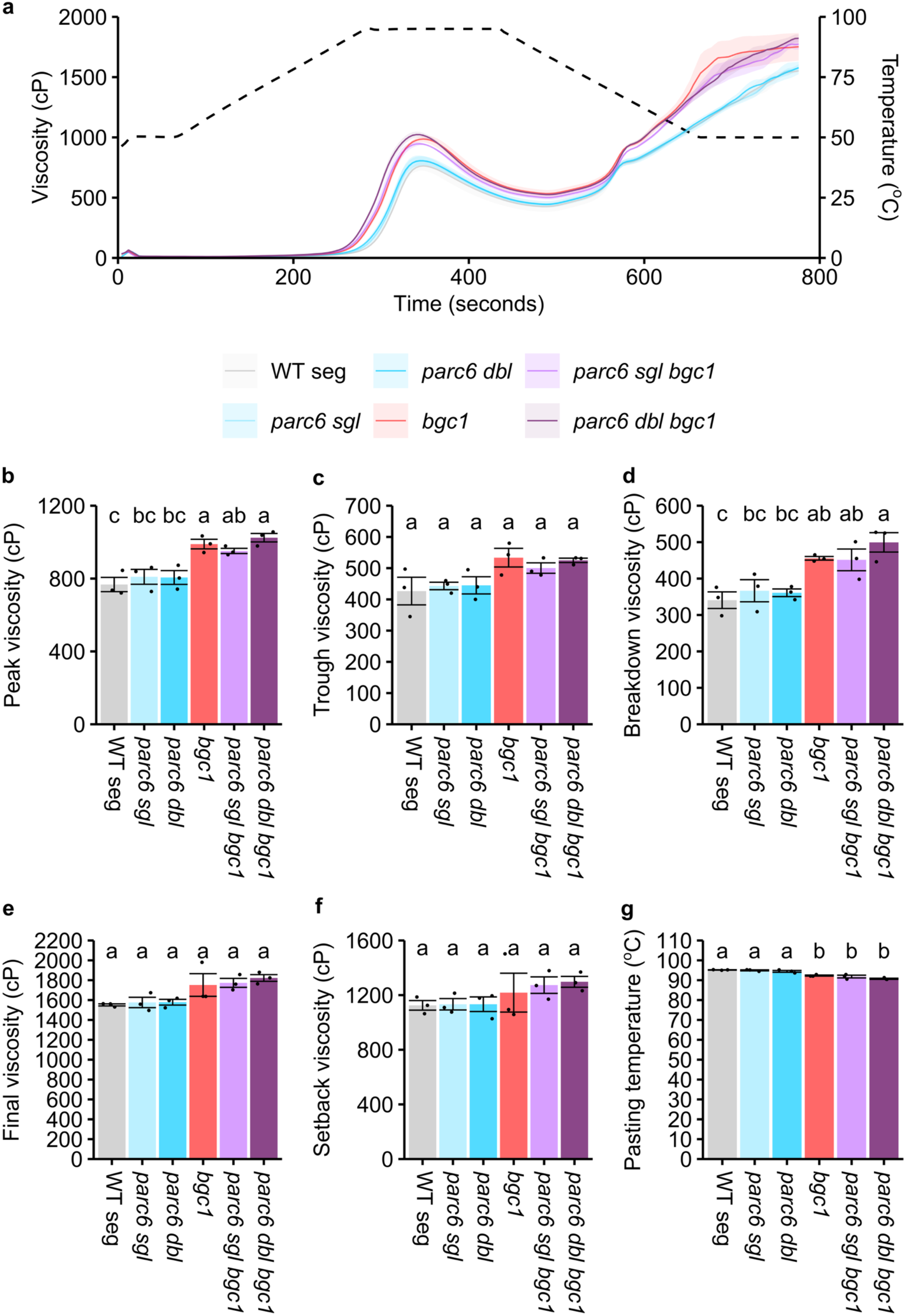
The *parc6 bgc1* mutants have increased peak and breakdown viscosities. (a) Rapid Visco Analyser (RVA) analysis of viscosity of purified starch (2 g in 25 mL water) during gelatinisation. Data are presented as means (solid line) ± SEM (shading) of three biological replicates, each using starch from grain harvested from a different field plot. Temperature changes are displayed as a dotted line on the right hand axis. Parameters from the viscographs in (a) were obtained. (b) Peak viscosity, (c) trough viscosity, (d) breakdown viscosity, (e) final viscosity, (f) setback viscosity, (g) pasting temperature. Data are presented as means ± standard error of the mean, with individual data points shown as black dots. Values with different letters are significantly different under a one-way ANOVA and Tukey’s post hoc test at *P* < 0.05 (*N* = 3 per genotype).

**Fig S7.**
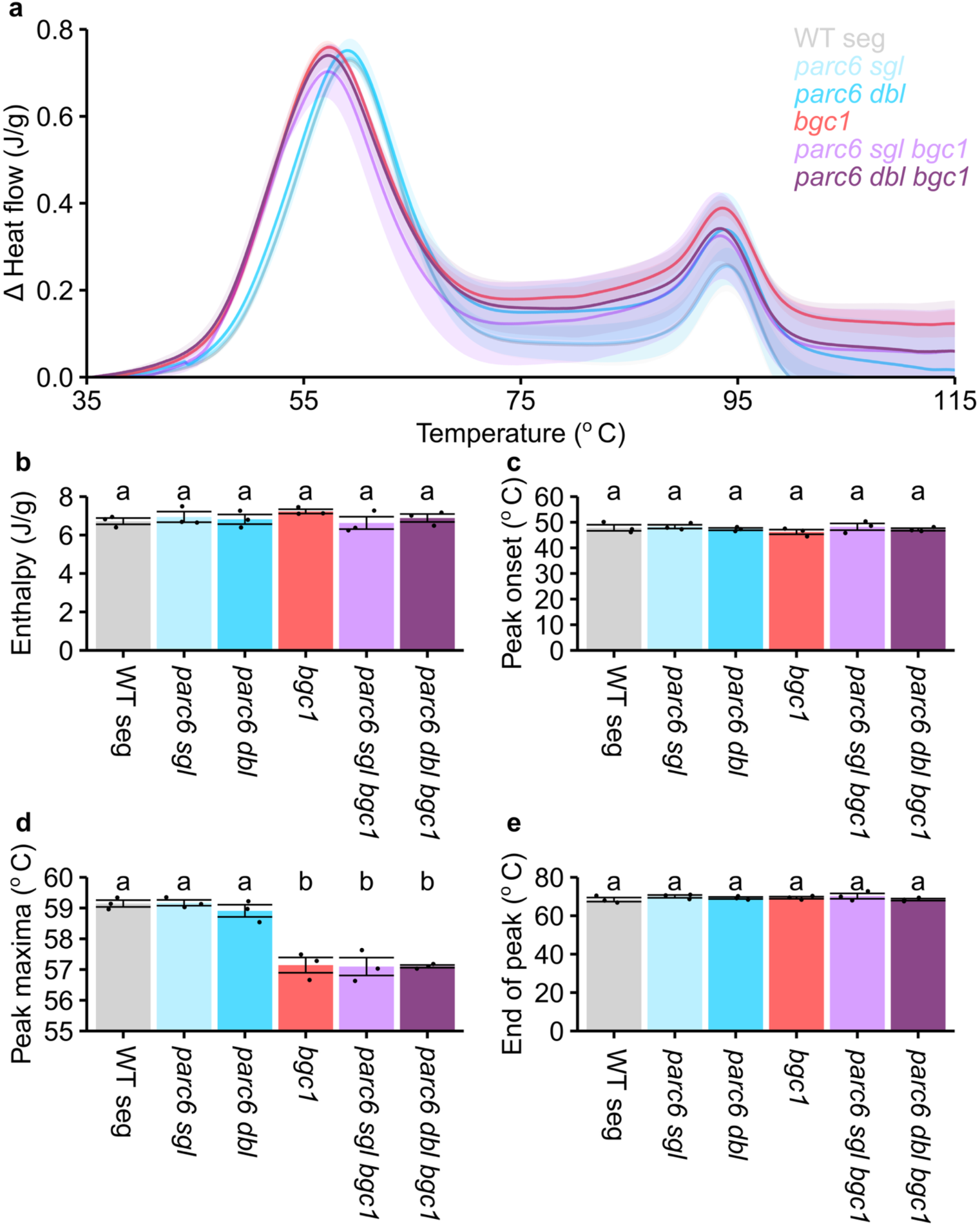
The *bgc1* starch has a shifted gelatinisation peak in Differential Scanning Calorimetry (DSC). (a) Purified starch (100 mg) was analysed on a DSC heating from 10°C to 150°C. The baseline heat flow was subtracted and data are presented as means (solid line) ± SEM (shading) of three biological replicates, each using starch from grain harvested from a different field plot. Only data from 35°C to 115°C are shown as this is where the peaks occur. The first peak was analysed - (b) enthalpy, (c) peak onset, (d) peak maxima, (e) end of peak. Data are presented as means ± SEM, with individual data points shown as black dots. Values with different letters are significantly different under a one-way ANOVA and Tukey’s post hoc test at *P* < 0.05 (*N* = 3 per genotype).

**Fig S8.**
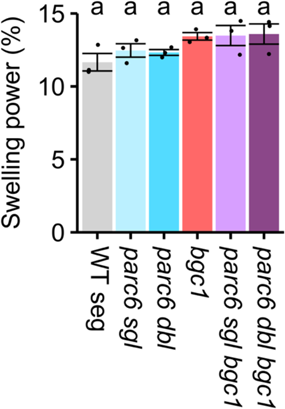
Starch from *parc6 bgc1* lines have no difference in swelling power. Swelling power of purified starch (10 mg) at 80°C. Data are presented as means ± SEM, with individual data points shown as black dots. Values with different letters are significantly different under a one-way ANOVA and Tukey’s post hoc test at *P* < 0.05 (*N* = 3 per genotype).

**Fig S9.**
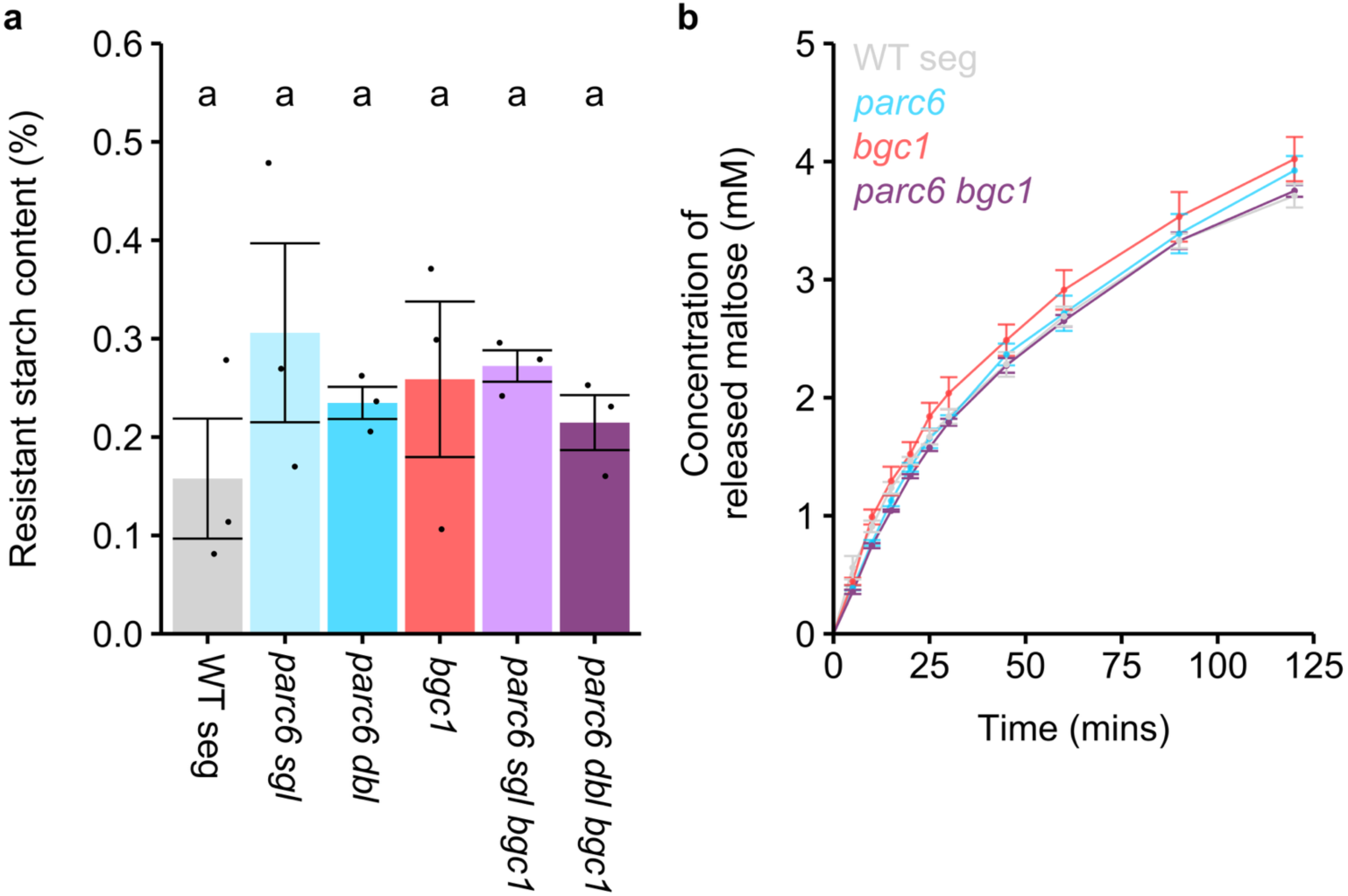
Purified *parc6 bgc1* starches have no differences in *in vitro* digestibility. (a) Resistant starch content of wholemeal flour. Data are presented as means ± SEM, with individual data points shown as black dots. Values with different letters are significantly different under a one-way ANOVA and Tukey’s post hoc test at *P* < 0.05. (b) Digestion of purified starch (100 mg) from WT seg, *parc6 dbl*, *bgc1* and *parc6 dbl bgc1* with pancreatic α-amylase. Data are presented as means ± SEM with values from three independent experiments, each using starch from grain harvested from a different field plot (*N* = 3 per genotype).

## Notes

### Competing Interest Statement

We declare that two of the authors (Lara Esch and David Seung) are co-inventors of a patent application describing our novel approach to increase granule size (PCT/EP2024/059060). We have no other conflicts of interest to declare.

